# High-throughput 3D tracking reveals the importance of near wall swimming and initial attachment behaviors of P. aeruginosa for biofilm formation on a vertical wall

**DOI:** 10.1101/2020.05.03.075507

**Authors:** Nicole Zi-Jia Khong, Yukai Zeng, Soak-Kuan Lai, Cheng-Gee Koh, Zhao-Xun Liang, Keng-Hwee Chiam, Hoi-Yeung Li

## Abstract

Studying the swimming behaviour of bacteria in 3 dimensions (3D) allows us to understand critical biological processes, such as biofilm formation. It is still unclear how near wall swimming behaviour may regulate the initial attachment and biofilm formation. It is challenging to address this as visualizing the movement of bacteria with reasonable spatial and temporal resolution in a high-throughput manner is technically difficult. Here, we compared the near wall (vertical) swimming behaviour of *P. aeruginosa* (PAO1) and its mutants Δ*dipA* (reduced in swarming motility and increased in biofilm formation) and Δ*fimX* (deficient in twitching motility and reduced in biofilm formation) using our new imaging technique based on light sheet microscopy. We found that *P. aeruginosa* (PAO1) increases its speed and changes its swimming angle drastically when it gets closer to a wall. In contrast, Δ*dipA* mutant moves toward the wall with steady speed without changing of swimming angle. The near wall behavior of Δ*dipA* allows it to be more effective to interact with the wall or wall-attached cells, thus leading to more capture events and a larger biofilm volume during initial attachment when compared with PAO1. Furthermore, we found that Δ*fimX* has a similar near wall swimming behavior as PAO1, however, it has a higher dispersal frequency and smaller biofilm formation when compared with PAO1 which can be explained by its poor twitching motility. Together, we propose that near wall swimming behavior of *P. aeruginosa* plays an important role in the regulation of initial attachment and biofilm formation.

**Importance:** Bacterial biofilm is a community of bacteria on surfaces which leads to serious problems in medical devices, food industry, and aquaculture. The initial attachment and subsequent microcolony formation play critical roles in bacterial biofilm formation. However, it is unclear how the initial attachment is regulated, in particular, on a vertical surface. To study this, we have developed a novel imaging technique based on light sheet microscopy, which overcame the limitations of other imaging techniques, to understand how 3D bacterial motility near a wall may regulate initial attachment during biofilm formation. Using our technique, we discovered that near wall swimming behavior of the bacteria, *P. aeruginosa*, plays an important role in the regulation of biofilm formation during initial attachment.

## Introduction

In the early stages of biofilm formation, planktonic bacteria swim close to the surface by rotating their flagella and attach to the surface using their pili. However, little is known about the dynamics of these processes (1). Several *P. aeruginosa* mutants have earlier been identified to be important in motility and biofilm formation (2). The mutant *ΔdipA* is found to have diminished swimming motility but enhanced initial attachment and reduced dispersal during early biofilm formation. *P. aeruginosa* lacking in DipA (*ΔdipA*) has significantly higher c-di-GMP levels, resulting in more efficient biofilm formation (3). In addition to flagella-mediated motility, *P. aeruginosa* is able to propagate at surfaces by type IV pili-mediated twitching motility. The type IV pili-mediated twitching motility contributes to the initial tethering and attachment during biofilm formation (4, 5). Mutant PAO1 lacking in the c-di-GMP binding protein, FimX, required for T4P assembly biofilm formation is less capable of microcolony and biofilm formation due to deficiency in twitching motility (6). Therefore, to fully understand the complex nature of biofilm formation, it is necessary to look into how each component of bacteria motility patterns contributes to the entire process of biofilm development.

Numerous motility strategies have been developed by bacteria to allow them to transverse complex natural environments and to facilitate cell-cell interactions (7, 8). By studying how bacteria move and analyzing their trajectories, we can extract valuable information on various microbial processes, such as behavioral responses towards chemical stimuli, signaling pathway mechanisms, as well as the behavioral signatures of different bacterial species during initial attachment of biofilm formation. However, it is extremely challenging to visualize and track bacterial movement and trajectories due to its small size and its variable dynamics, which can span a broad range from milliseconds to minutes (9). The standard approach to examine bacterial motility is to carry out two-dimensional (2D) imaging using optical microscopy (10, 11) or perform swarming assays on agar plates (12). Nevertheless, as most microbial systems are intrinsically three-dimensional (3D) in their organization, 2D approaches may lead to misinterpretation of behavioral patterns.

Several techniques have been developed to track 3D motility behavior of bacteria. One of the earliest techniques was based on the automatic motion of the scanning stage to keep an individual bacterium in focus (13). This technique is limited to observing one cell at a time but provided key understanding of the motility behavior of *Escherichia coli*. More recent and powerful optical techniques that have been used include intensity correlation microscopy, defocused microscopy (14), stereoscopic microscopy (15) and digital holographic microscopy (15–17). Digital holographic microscopy, in particular, is a preferred method to perform 3D tracking of bacteria because it is able to capture a large depth of field and therefore providing high throughput data. However, it is often limited by the lack of contrast of the samples (especially when using unlabeled cells) and secondary scattering of the bacteria, which produce poor holograms due to overlapping signals when there are too many bacteria cells (16). These make resolving and tracking of individual cell difficult and more errors are generated when the concentration of the sample is high. Although these techniques aim to image 3D motility of bacteria, they are not able to analyze biofilm formation on vertical surfaces.

Here, we introduce a novel imaging technique that is sensitive, minimally toxic and can rapidly capture large fields of view for 3D measurement of individual motile bacterium using the light sheet microscopy. The 3D trajectories obtained do not suffer from the errors mentioned above. Furthermore, our imaging technique allows proper visualization of bacteria behavior near a vertical wall. Biofilms commonly form on various solid surfaces, including vertical and tilted walls, ceilings and pipes (18).To our knowledge, many studies focused on biofilms formed on air-water interface or on a horizontal surface (19). The imaging technique described in this paper takes into consideration bacteria motility near and away from a vertical wall, representative of naturally occurring biofilms. By using fluorescence-labeled cells, we can directly visualize and track multiple individual cell trajectories and obtain information about swimming speeds and turns. The design of our technique also allows us to capture swimming patterns of bacteria in bulk fluid and near wall surfaces. This robust method provides the means to study the dynamics of bacterial motility and also allows us to examine the structural architecture and microbial processes of microbial communities on a wall. By comparing the free swimming, near wall dynamic behaviors and initial attachment on a wall of *P. aeruginosa* (PAO1) and PAO mutants Δ*dipA* and *ΔfimX*, we are able to understand the correlation between near wall behavior and initial attachment in the regulation of biofilm formation.

## Results

### 3-Dimensional light sheet microscopy system setup and design for tracking of multiple individual bacterial cell trajectories

To visualize the swimming behavior of individual bacterial cells using light sheet microscopy, we constructed chambers loaded with fluorescently labelled cells and imaged their swimming behavior over a defined time course. The chamber was made of 1% Luria-bertani (LB) agarose to support biofilm formation. PAO1, *ΔdipA or ΔfimX* were first incubated in the chamber with ABTGC minimal medium for 6 hours to allow initial attachment. The agarose chamber was then mounted in the Zeiss Z1 light sheet microscope for imaging. (Fig. 1A). By moving the sample into the z-direction, the trajectories of bacterial cells in 3D can be obtained. During light sheet image acquisition in the z-direction continuously, the x-y coordinates of the bacterial cells in each z-step can be captured while the exposure time (i.e 80 ms) of each z-step are recorded as time intervals (Fig. 1B). If the bacteria turn and swim in the reverse direction during image acquisition, the trajectory will be cut off with shorter time durations (Fig. 1C). However, owing to rotational diffusion, the bacterial cell will typically not execute a sharp turn but will instead exhibit a “curved” trajectory during reversal. Part of this curved trajectory that is moving forwards can still be captured and hence we can still capture and record a reversal taking place. The x-y coordinates of bacterial cells in each z-step image are linked and the acquisition time of each z-step is taken into account to reconstruct the trajectories (Fig.1D). When all information is combined, a space-time projection can be created, and we are able to determine the bacterial trajectories in 3D over a period of time. The space-time projection and the 3D trajectories of individual bacterial cells in a large field of view can easily be visualized by a simple 3D reconstruction algorithm (Fig. 1E). We are able to track individual bacterial cells for a duration of 15 s. In general, we can track hundreds of cells simultaneously to obtain reliable statistics for swimming speeds and turns.

**Figure 1.**
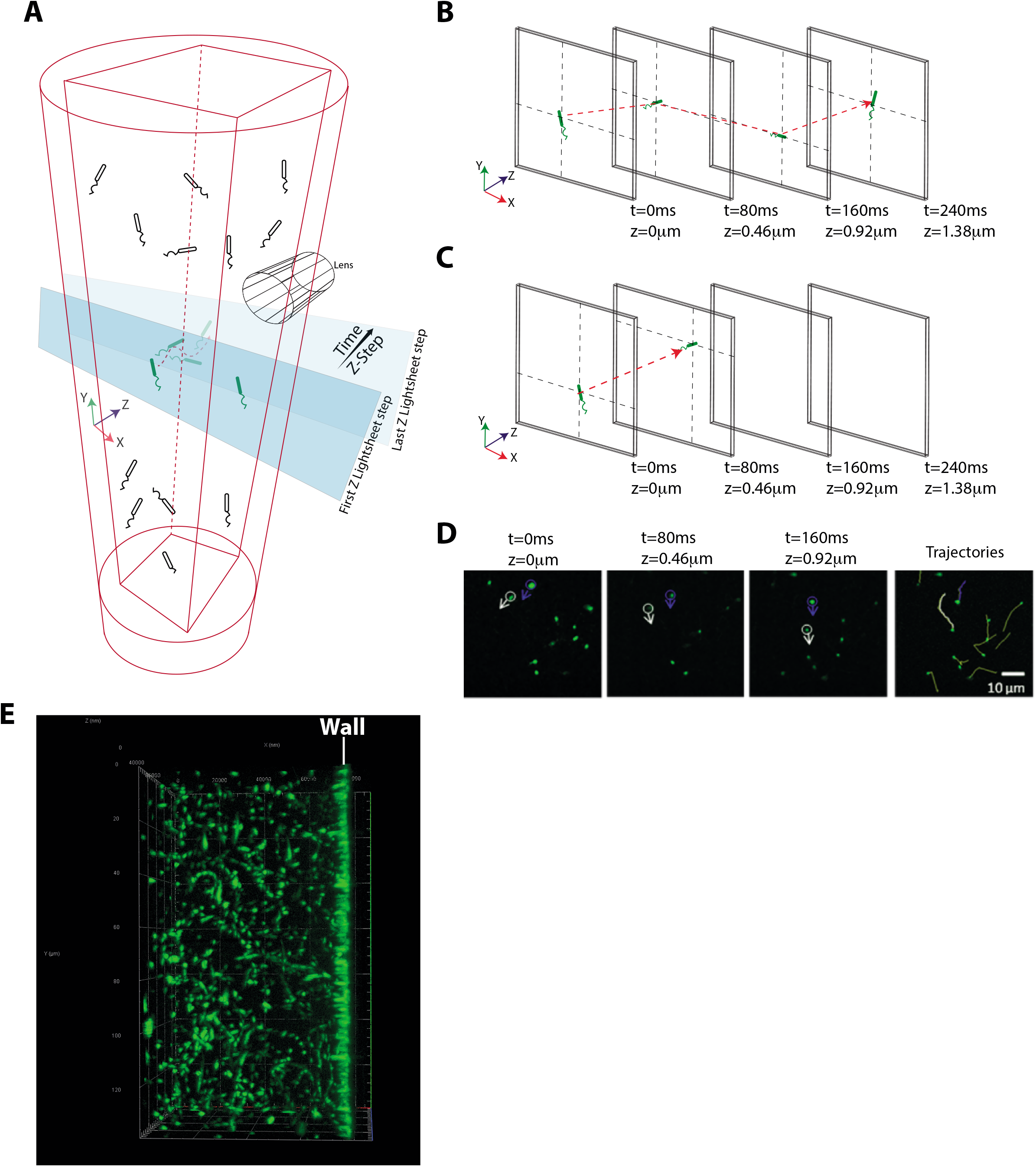
Illustration of light sheet microscopy system setup and design. (A) Fluorescence-labeled bacterial cells were loaded into custom made agarose chamber (1% agarose in ABTGC medium that supports biofilm formation). The illumination beam path is orthogonal to the detection beam path. A thin sheet of light is formed at the focal plane of the detection objective. The sample is moved in the z-direction immediately after each acquisition and 3D image of the sample is reconstructed thereafter. (B) During light sheet imaging acquisition in the z-direction, the x-y coordinates of bacterial cells in each z-step can be captured while the exposure time (i.e 80 ms) of each z-step can be used as time intervals. (C) During light sheet image acquisition, the trajectory will be cut off with shorter time durations when bacteria turn and swim in a reverse direction. (D) Tracking of bacterial cells from the frame by frame analysis. Bacterial cells which appear as “particles” are stitched together to form trajectories using the linear assignment problem (LAP) method after segmentation utilizing the difference of Gaussians approach and filtering based on mean intensity and quality. (E) Large field of view of the 3D trajectories of bacteria swimming in the agarose chamber.

We incubated PAO1, *ΔdipA* and *ΔfimX* mutants in the ABTGC medium within the chamber described in Figure 1A, and light sheet imaging was carried out after 6 hours of incubation. We found that, in the bulk medium and away from walls, PAO1,*ΔdipA* and *ΔfimX* swam with an average speed of 23.9±6.0 μm/s, 23.6±3.9 μm/s and 24.2±5.0 μm/s, respectively (Fig 2A & B & Table 2). These values are consistent with previously reported values (2). The average speeds were obtained by averaging the speeds of individual cells over their whole tracked trajectories. To further validate our imaging and tracking method, we repeated the experiments with *E. coli*, we found that *E. coli* cells swam with an average speed of 15.5±3.9 μm/s (Fig. 2B, S1A & Table 2), also consistent with previously reported values (20).

**Figure 2.**
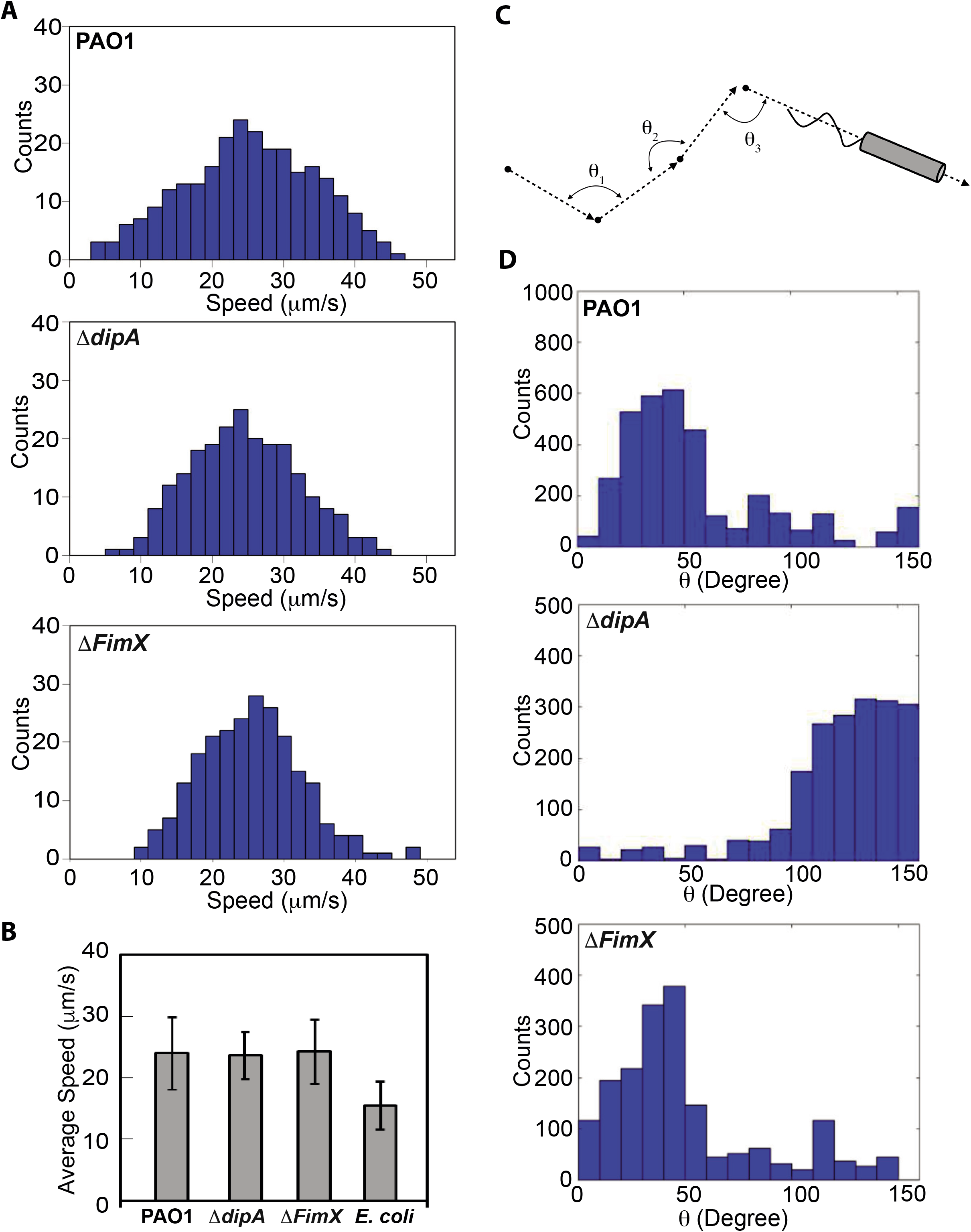
Swimming behavior of PAO1,*ΔdipA* and *ΔfimX*. (A) Histograms of the speed of *PAO1, ΔdipA and ΔfimX* obtained from 323, 598, and 375 cell trajectories tracked from 5 different experiments, respectively. (B) Average speeds for *PAO1, ΔdipA, ΔfimX and E. coli* are 23.9 ±6.0 μm/s, 23.6 ±3.9 μm/s, 24.2 ±5.0 μm/s, and 15.5 ±3.9 μm/s, respectively. (C) Turning angles for PAO1, *ΔdipA, ΔfimX and E. coli*. The turning angle, θ, represents the angle between stitched line segments between two adjacent frames. It is the angle between two straight lines, e.g., θ_1_ is the angle through two-point locations of the same tracked particle in frames i and i+1. (D) The distribution of turn angles θ during a swimming trajectory was quantified and analysed for PAO1 and *ΔfimX* and *ΔdipA*.

In addition, we can also track when individual bacterial cells change their swimming directions and “turn”. For every segment of a cell’s trajectory, we can define a turning angle, θ, as the angle between stitched line segments between two adjacent frames, i.e., it is the angle between two straight lines: the first drawn through the two-point locations of the particle in frames *i* and *i+1*, and the second drawn through the two-point locations of the particle in frames *i+1* and *i+2* (Fig. 2C). Unlike the more commonly studied “run-and-tumble” mechanism of E. coli, where counter-clockwise (CCW) rotation of the flagella leads to forward motion and clockwise (CW) rotation leads to tumble and hence change of direction, *P. aeruginosa* makes use of a different mechanism to swim [21], where CCW rotation leads to forward motion and CW rotation leads to backward motion. During these reversals from CCW to CW or vice versa, the flagellum typically is not rotating, in a phase which we term “pause.” During these pauses, rotational diffusion leads to the cell adopting a new direction. We term this mechanism “run-reverse-pause”.

For PAO1, we found that a typical trajectory comprised a series of straight tracks interspersed with sharp turns. These sharp turns suggest that monotrichous PAO1 indeed swim in a “run-reverse-turn” mechanism, where a run corresponds to the single helical flagellum rotating counter-clockwise and reverse to the flagellum rotating clockwise, with the sharp turns corresponding to changes in the direction of flagellum rotation (21). These results are shown in Fig. 2D, which shows a peaked distribution in the turning angles of PAO1 for 0 ≤ θ ≤ 50 degrees. In contrast, the distribution of turning angles for *ΔdipA* shows a peak at larger angles (θ ? 100 degrees), corresponding to curved trajectories with no sharp turns. Thus, *ΔdipA* mutants exhibit reduced turnings, i.e., reduced flagellar reversals. This is consistent with previous findings for dipA [34]. Furthermore, *ΔfimX* exhibited a similar turning angles to PAO1 for 0 ≤ θ ≤ 50 degrees. For *E. coli*, a broad distribution with no peaks is observed, where individual *E. coli* cells exhibit “tumbles” that are distributed over a wide range of angles, consistent with the “run-and-tumble” mode of *E. coli* swimming (Fig S1B).

### Near wall behavior of bacterial cell swimming

The presence of a wall can significantly modify the swimming behavior of a bacterial cell that is moving near it. Although there are many theoretical studies on the interactions between cells and boundaries (22), there are relatively fewer reports of experiments which quantitatively study cell swimming behavior near a wall. One plausible reason could be the difficulty associated with imaging and tracking only those cells that are close to the wall, which will require a three-dimensional setup. There have been several studies on three-dimensional tracking of individual bacterial cells (20, 23). However, these studies were not able to track the behavior of individual cells near wall with reasonable resolution.

Here, we focus on tracking individual cells that are swimming close to a wall. We measure quantitatively the changes in the swimming speed, trajectories, and swimming orientation that occur for bacterial cells swimming close to a wall.

### PAO1 and ΔfimX changes its swimming speed near a wall but not ΔdipA

The change in swimming speed near a wall has been addressed theoretically (24–26) and we shall not repeat the theoretical calculations here. Briefly, we expect the swimming speed to increase as the cell swims closer to the wall. Heuristically, this can be explained as follows. Near to a wall, the viscous drag experienced by a swimming bacterial cell increase. However, for a rod-like bacterial cell propelled by a rotating flagellum, the component of the drag coefficient in the direction perpendicular to the long axis of the rod increases faster than the parallel direction. Hence, the swimming speed which varies with the ratio of the perpendicular drag coefficient to parallel drag coefficient, increases. We plot the speeds of individual cells (μm/s) as a function of how far they are from a wall (*h*). If we fit how the speed *v* varies with the perpendicular distance to the wall *h* with the form *v ~ h^-b^*, we obtain the relation *β=* 0.13 for PAO1, 0.056 for *ΔdipA* and 0.26 for *ΔfimX*. (Fig 3A). This shows that the speed of *ΔdipA* does not increase as it swims near to the wall. Thus, PAO1 and *ΔfimX* cells indeed swim faster (average speed of 39.3 ± 6.2 μm/s and 36.4 ± 5.2 μm/s) when they are closer to the wall (*h* < 5 μm). However, for *ΔdipA*, this trend is not present; the mutant cells do not exhibit higher swimming speed (28.1 ± 6.2μm/s) when they are closer to the wall (Fig 3B and Table 2). We wish to reiterate that the swimming speed of PAO1, *ΔdipA and ΔfimX* far from the wall are similar (Fig. 2A & B and Table 2). Our finding suggests that the *ΔdipA* mutation affects the swimming speed only when the cells are near a wall, necessitating a technique to image and track them near a wall such as we are describing here. Similarly, *E. coli* exhibit a higher speed when they are approaching the wall (Fig S1A & 3B, Table 2)

**Figure 3.**
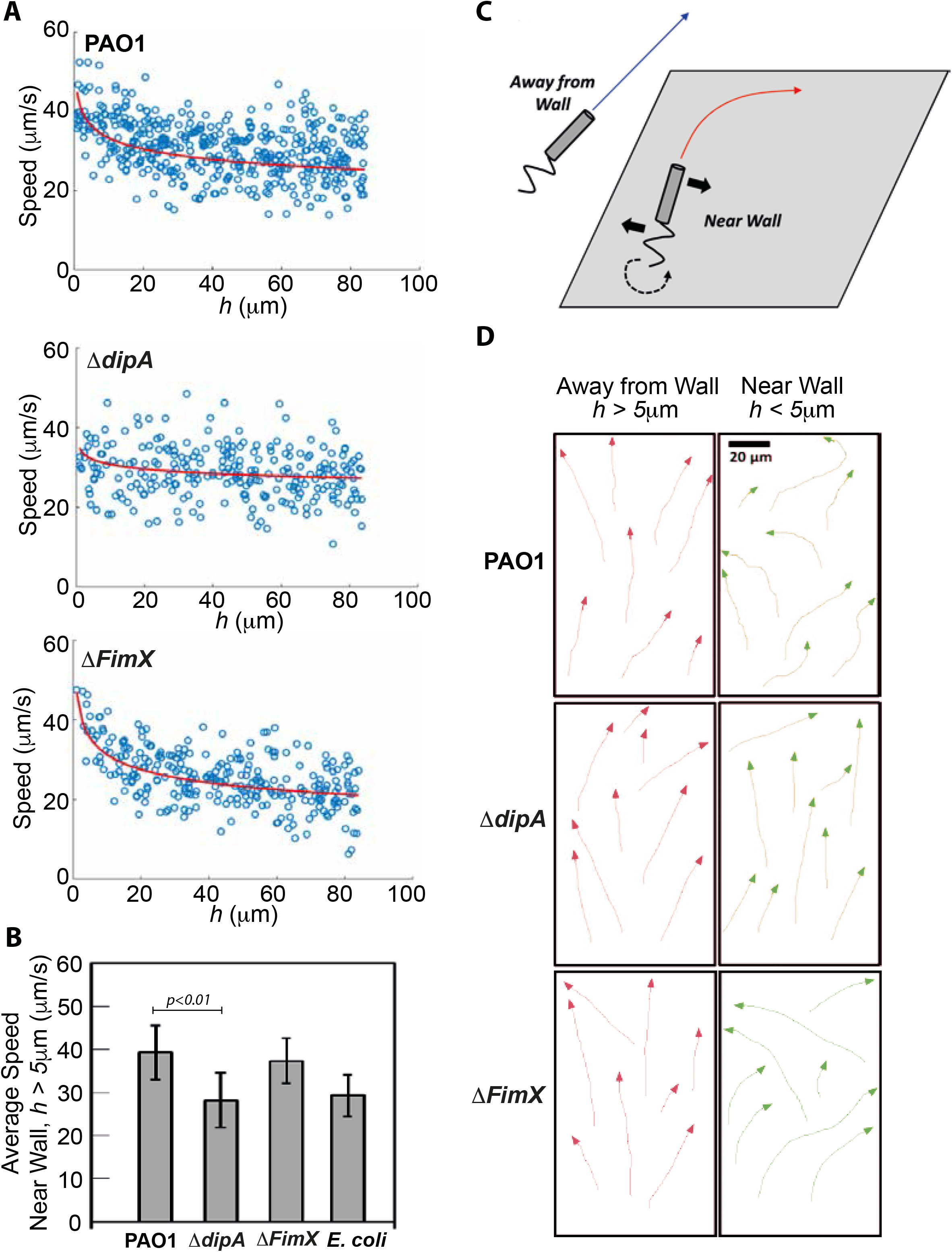
Near wall swimming behavior of PAO1, *ΔdipA and ΔfimX*. (A) Cell speed vs. *h* which is the perpendicular distance to the wall. The red line is a least-squares fit to the form *v ~ h^-β^* for some exponent *β*, where *β* = 0.13 for PAO1, 0.056 for *ΔdipA*, 0.26 for *ΔfimX*. (B) The histogram shows the average speeds near the wall (*h* < 5 μm) for PAO1*, ΔdipA and ΔfimX*. (C) Illustration of swimming trajectory of bacteria with helical flagella near a wall and away from a wall. A force perpendicular to the direction of motion and parallel to the wall is generated when helical flagella rotate. An equal and opposite force acts on the cell body and causes it to rotate in the opposite direction as the flagella (solid black arrows). This creates a net torque which results in the cell rotating (dashed black arrow). (D) Trajectories for PAO1, *ΔdipA and ΔfimX* away from a wall (*h* > 5μm) and near a wall (*h* < 5μm). Circular trajectories are observed near the wall of PAO1 and *ΔfimX*, but not away from the wall. *ΔdipA* shows trajectories with large radius of curvature (i.e., effectively straight) regardless of whether it is near (*h* < 5μm) or away from the wall (*h* > 5μm).

### PAO1 and ΔfimX cells change their trajectories, but not ΔdipA cells, when they are near a wall

For bacteria with helical flagella such as *E. coli*, they are known to change their trajectories from straight to circular when swimming near a wall (27, 28). When the helical flagella rotate, they generate a force that is perpendicular to the direction of motion and parallel to the wall. There is an equal and opposite force acting on the cell body, which causes it to rotate in the opposite direction as the flagella. Thus, there is a net torque which results in the cell rotating (dashed black arrow in Fig. 3C). The flagella of *P. aeruginosa* cells rotate in both counter-clockwise (for “running”) and clockwise (for “reversing”) resulting in the cells turning both to the right and left. Trajectories of PAO1, *ΔdipA* and *ΔfimX*cells away from a wall and near a wall are shown in Fig. 3D. Interestingly, the radius of curvature of the trajectories of *ΔdipA* mutants are larger, i.e., their trajectories appear “more straight” than those of PAO1. This could be due to the slower swimming speed and rotation rate of the mutant cells (22). Furthermore, we also see right-handed turns only for *E. coli* when they moved closer to the wall (Fig. S1D). Since the flagella of *E. coli* cells rotate in the counter-clockwise direction for propulsion, the cells constantly turn only to the right.

### PAO1 and ΔfimX cells change their orientation, but not ΔdipA cells, when they are near a wall

As a cell moves towards a wall, it will be reoriented to become parallel to the surface of the wall (11). Referring to the sketch in Fig. 4A, if a cell is moving towards a wall at an orientation φ (dashed red arrow), then there will be a gradient in the flow field that will cause the cell to rotate until φ =0 (dashed blue arrow). The cell trajectory angle of approach towards the wall, φ, is obtained for each individual cell trajectory by finding the angle between two lines: the first line is drawn through the location of the particle in the first frame and the point on the wall closest to it, while the second line is drawn through the location of the particle in the first and last frame. As such, φ = 0 and φ = 90° denote cells with trajectory paths running perpendicular and parallel to the wall, respectively. In Fig. 4B, we plot the distribution of orientation φ as a function of distance *h* from the wall. For both PAO1 and *ΔfimX*, we see that the orientation φ approaches 90 degrees as *h* decreases, consistent with the expectation that the cells reorient themselves to be parallel to the wall (Fig 4B). Similarly, *E. coli* exhibit the same behaviour to reorient themselves to be parallel to the wall (Fig. S1E). However, *ΔdipA* cells do not reorient themselves (Fig 4B). Taken together, our observations suggest that *ΔdipA* cells do not increase their speed nor reorient to become parallel to the wall as they approach a wall.

**Figure 4.**
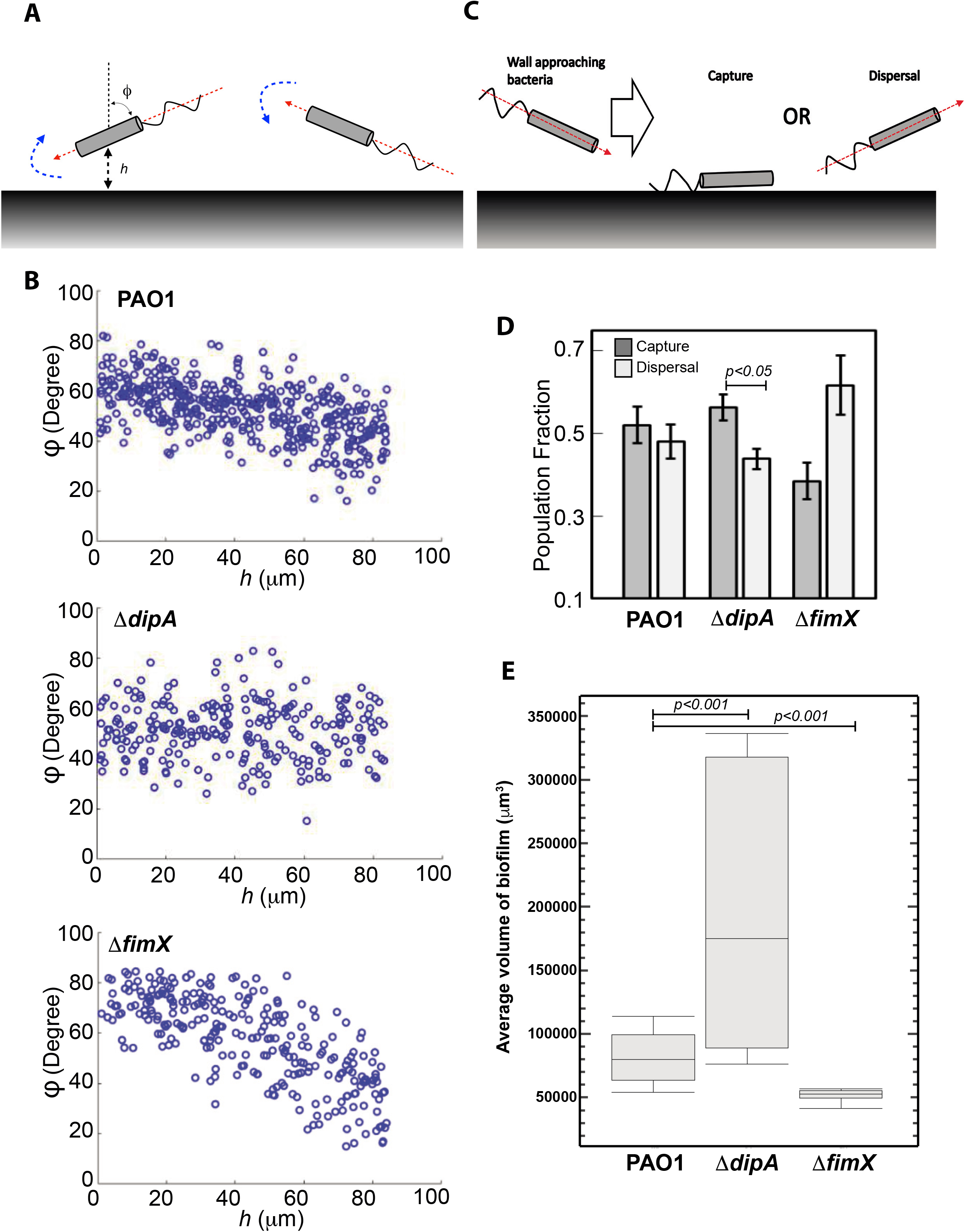
*ΔdipA* does not reorient to be parallel to the wall surface, which correlates to a higher capture frequency for better biofilm formation. (A) Swimming bacteria reorient to be parallel to the surface of the wall. When a cell moves towards a wall at an orientation φ (dashed red arrow), there will be a gradient in the flow field that will cause the cell to rotate until φ =0 (dashed blue arrow). (B) The cell trajectory angle of approach towards the wall, φ, is obtained for each individual cell trajectory. The number of PAO1 and *ΔfimX* with φ ending to 0 increases as the distance to the wall *h* decreases. (C) Schematic of bacterial cell approaching a wall, capture and dispersal. (D) *ΔdipA* showed a significant higher percentage of capture event than PAO1 and *ΔfimX* whereas *ΔfimX* showed a significant higher percentage of dispersal when compared with PAO1 and*ΔdipA*. (E) The average volume of biofilm of PAO1, *ΔdipA and ΔfimX*.

### *ΔdipA* cells forms larger biofilm because of the intrinsic near wall swimming behavior whereas *ΔfimX* cells forms smaller biofilm because of high dispersal frequency

It is important to understand how near wall behaviour could correlate to the capture and dispersal events during initial attachment stages of biofilm formation. We first analysed the light sheet images by measuring the population fraction of cells that are captured versus those that are dispersed, for both PAO1, *ΔdipA* and *ΔfimX* (Fig. 4C). We consider a cell to be captured when the average of its near-wall velocity vector is pointing towards the wall, and dispersed when it is pointing away from the wall. We found that 52% population of PAO1, 56% population of *ΔdipA* and 39% population of *ΔfimX* were captured, respectively; and 48% population of PAO1, 44% population of *ΔdipA* and 61% population of *ΔfimX* were undergoing dispersal, respectively (p-value <0.05) (Fig 4D). In addition, we measured the biofilm volume (biomass) on a wall surface with area of 23,267 μm^2^ using 3D rendering images (Imaris) to show the correlation between near wall behavior and biofilm size. We found that *ΔdipA* formed a largest volume of biofilm whereas *ΔfimX formed a smaller biofilm* after 6 hours of incubation (Fig. 4E). These are consistent with other reports, which suggest that *ΔfimX* is deficient in biofilm formation while *ΔdipA* demonstrated an enhanced initial attachment and lower dispersal during biofilm formation (3, 6).

## Discussion

The study of the spatial dynamics of bacteria is crucial to formulate solutions against microbial infections and biofouling. It is challenging to develop an imaging system, which is fast enough to observe the spatiotemporal activities of free-swimming bacteria with sufficient resolution. We have developed a new technique based on light sheet microscopy to observe single bacterium swimming behavior with better precision and resolution compared to other pseudo-3D imaging methods such as holographic particle tracking. Consequently, we are able to accurately measure bacterial swimming velocity and trajectories that are consistent with existing theoretical predictions and experimental data. It is known that bacterial swimming behavior is largely influenced by the presence of solid surfaces and the swimming behavior near a wall is significantly different compared to ‘away from wall’ regions (29).

Pathogenic microorganisms colonize and form biofilms on various solid surfaces, causing severe environmental damage and pollution(18). In our light sheet microscopy imaging setup, we built a chamber that allows concurrent and clear imaging of bacteria movement near a vertical wall and in regions away from the wall (Fig. 1A). Our method is suitable for studying the behaviour of bacteria motility on walls and tilted surfaces, which contributes to understanding and then tackling the problem of biofouling of ships hulls and pipelines.

Using the technique described in this paper, we investigated the swimming behavior of *P. aeruginosa* and its mutants that exhibit varied swimming patterns and ability of biofilm formation. We then imaged and quantified the statistics of trajectories, speed, and orientation, which can provide valuable insights into bacterial behavior in both ‘near wall’ and ‘away from wall’ environments. This technique is suitable for characterizing flagella-dependent motility by comparing wild type species to mutants with defective flagella movement of other bacterial species to understand mechanistically the function of various bacterial genes implicated in biofilm formation.

Future applications of light sheet microscopy to visualize bacterial swimming dynamics are vast, especially to naturally occurring bacterial communities. One useful application is to image the initial phases of bacterial biofilm formation in 4D, the fourth dimension being time. To date, it remains challenging to observe the early stages of biofilm formation in 4D due to obstacles in imaging techniques. The speed of bacterial displacement and attachment to early biofilm layers are technically difficult to capture without using our approach. Furthermore, the use of different fluorescence labels makes it possible to image more than one population or species of bacteria concurrently, within the same field of view. One could couple spatiotemporal analyses of bacterial swimming with the chemotactic response to varying chemo-effectors to understand the influence of nutrients in early biofilm formation. The first step to treating biofouling is to understand early-stage bacterial dynamics through direct imaging of bacteria-surface interactions. Our method developed and described here is robust and applicable to other living microbial systems. Therefore, we envisage that this technique will transform modern 3D microscopy with potentially massive practical capabilities.

Our findings suggest that PAO1 increases its speed and change its swimming angle when it gets closer to a wall. This will result in less capturing and more dispersal events, which eventually results in smaller biofilm size during initial attachment. The motility behaviour of the pilus deficient FimX mutant, which is less adept in bacterial colonization of surfaces and formation of biofilms, formed biofilms indistinguishable from those of WT PAO1. In contrast, Δ*dipA* mutant moves toward the wall with steady speed without changing of swimming angle. The near wall behavior of Δ*dipA* allows it to interact more effectively with the surface or other bacteria leading to more capturing and less dispersal events. Thus, a larger biofilm can be formed by Δ*dipA* during initial attachment.

## Materials and Methods

### Bacteria strains and culture

All bacteria strains used in this study is listed in Table 1 in the supplementary material. All bacteria were grown in LB broth overnight with agitation at 37°C. Before imaging, the bacteria were diluted to an optical density of OD_600_=1.5 and stained with Vybrant^®^ Dyecycle Green^™^ stain (Invitrogen) for 30 minutes under agitation.

**Table 1.**

| Strain | Characteristic | Source/Reference |
| --- | --- | --- |
| *E. coli* DH5α | Competent cells used for molecular cloning | Invitrogen |
| *P. Aeruginosa* PAO1 | Wild type strain from the Washington Genome Center PAO1 mutant library | (30) |
| $\Delta dipA$ | PA5017 transposon mutant PW9424 from the Washington Genome Center PAO1 mutant library | (30) |
| $\Delta fimX$ | PA4959 transposon mutant PW9347 obtained from the Washington Genome Center PAO1 mutant library | (30) |

**Table 2.**

| | Mean speed ($\mu\text{m/s}$) | | P-value (Away from wall vs Near wall) |
| --- | --- | --- | --- |
| | Away from Wall ($h > 5\mu\text{m}$) | Near Wall ($h < 5\mu\text{m}$) | |
| PAO1 | $23.9 \pm 6.0$ | $39.3 \pm 6.2$ | $p < 0.01$ |
| $\Delta dipA$ | $23.6 \pm 3.9$ | $28.1 \pm 6.2$ | $p > 0.05$ |
| $\Delta fimX$ | $24.2 \pm 5.0$ | $36.4 \pm 5.2$ | $p < 0.01$ |
| *E. coli* | $15.5 \pm 3.9$ | $29.2 \pm 4.9$ | $p < 0.01$ |

### Light sheet microscopy imaging

The light sheet microscopy imaging used in this study requires a custom-made apparatus that included a bacteria inoculation chamber and Zeiss light sheet Z.1 microscope (Carl Zeiss). The bacteria inoculation chamber was made of 1.5% LB agarose. For biofilm formation, the stained bacteria was pelleted down, resuspended in specially formulated ABT minimal medium supplemented with 2 g of glucose per litre and 2 g of Casamino Acids per litre (ABTGC) (31) and loaded into the agarose chamber. After 6 hours of incubation at 37°C, the agarose chamber was mounted onto the Zeiss light sheet Z.1 microscope. The 3-dimensional imaging boundary was defined within 200 z-steps along the z-axis with 0.46μm thickness. Each z-step was recorded at an exposure time of 80 ms using a 40x water immersion objective (N.A 1.0) and a high-speed camera (Carl Zeiss). The z-step moved to the next z-step immediately after each acquisition without any delay (Fig 1A). Each experiment was repeated for 60 times in three independent experiments.

### Data analysis

The trajectories of individual bacterial cells are obtained from the frame by frame analysis of the captured images. In a single frame, individual bacterial cells appear as “particles.” Automated particle tracking is then used to stitch together the “particles” to form trajectories. Algorithms for particle tracking have been extensively reported. Briefly, the particles are first segmented and identified using the difference of Gaussians approach (32). They are then filtered based on mean intensity and quality. Next, the segmented particles are tracked by linking the individual particles from each frame to the next using the linear assignment problem (LAP) method (33), with modifications to the linking cost calculations with respect to both mean intensity and quality. An example of this tracking is shown in Fig. 1D.

## Acknowledgments

This project is supported by Singapore Ministry of Education Academic Research Funding Tier 1 (RG44-16) and Nanyang Technological University SUG to H.Y.L. and Tier 1 (RG138/16) to C.G.K.

