## Supplementary Figure 1 for "High-throughput 3D tracking reveals the importance of near wall swimming and initial attachment behaviors of P. aeruginosa for biofilm formation on a vertical wall"

### Figure S1

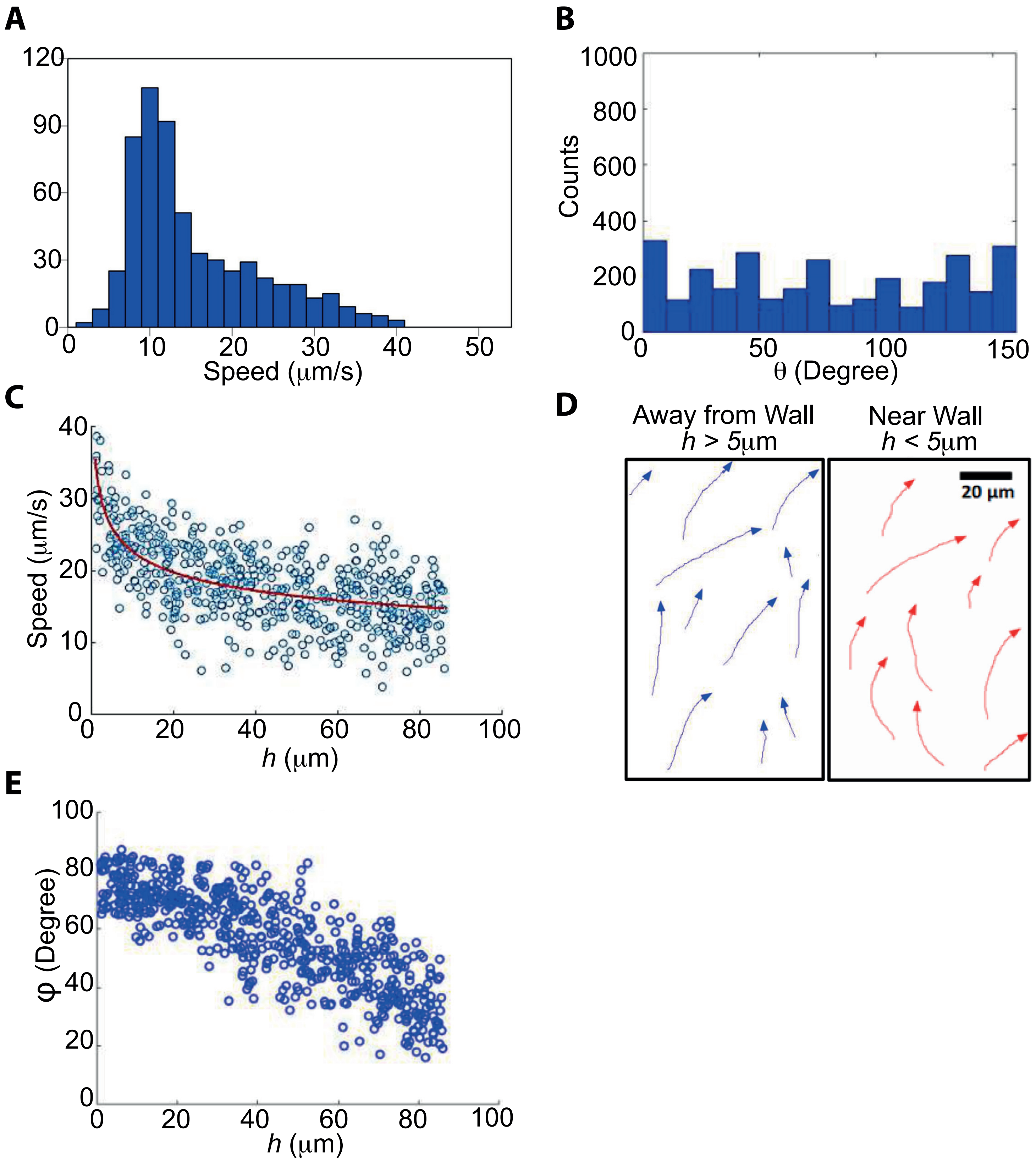

A
120
90
60
30
0
0
10
20
30
40
50
Speed (μm/s)
B
1000
800
600
400
200
0
0
50
100
150
Counts
θ (Degree)
C
40
30
20
10
0
0
20
40
60
80
100
Speed (μm/s)
h (μm)
D
Away from Wall
h > 5μm
Near Wall
h < 5μm
20 μm
E
100
80
60
40
20
0
0
20
40
60
80
100
φ (Degree)
h (μm)

Figure 1S. Near wall swimming behavior of *E. coli*. (A) Histograms of the speed of *E. coli*. (B) The turning angle, θ, for *E. coli* showed a broad distribution throughout.
(C) *E. coli* speed vs. *h* which is the perpendicular distance to the wall. The red line is a least-squares fit to the form $v \sim h^{-\beta}$ for some exponent $\beta$, where $\beta = 0.12$ for *E. coli*. (D) *E. coli* shows right-handed turns only when it swam near a wall (E) The cell trajectory angle of approach towards the wall, φ, is obtained for each individual cell trajectory. The number of *E. coli* with φ ending to 0 increases as the distance to the wall *h* decreases.
